# ASCL1 and OLIG2 Expression Dynamics Control Glial Cell Fate and Regional Diversity in the Dorsal Forebrain

**DOI:** 10.64898/2026.02.02.703158

**Authors:** Luis E. Paez-Beltran, Haojie Chen, Milindu Liyanapathirana, Estrella Villicana, Bianca L. Myers, Jinyi Duan, Antonella Riega, Tou Yia Vue

## Abstract

Gliogenesis is a multistep process that begins with the specification of glial progenitors (GPs) into migrating and proliferating precursor cells, which later differentiate into mature astrocytes and oligodendrocytes. How these developmental processes are coordinated to generate the diverse glial lineages in gray matter (GM) and white matter (WM) in the brain remains poorly understood. Here, we show that the basic-helix-loop-helix (bHLH) transcription factor ASCL1 serves as a direct mechanistic link between glial cell fate specification, migration, proliferation, and differentiation in the dorsal forebrain. Notably, ASCL1 is dynamically expressed in GPs, initiating in the ventricular zone (VZ), peaking in the intermediate zone (IZ), but is downregulated once GPs enter the cortical plate. Lineage tracing of ASCL1+ GPs demonstrates that they subsequently co-express OLIG2 to generate both astrocytes and oligodendrocytes in an “outside-in” pattern starting from the upper cortex inward to the corpus callosum, the opposite pattern of neurogenesis. Gain- and loss-of-function experiments further reveal that a sustained ASCL1 expression is essential for inducing sufficient levels of OLIG2 required to specify oligodendrocyte precursor cell (OPC) fate. Interestingly, a persistent ASCL1 expression also maintains OPCs into postnatal stages by promoting their self-renewal while suppressing their differentiation into postmitotic oligodendrocytes. Together, these findings establish ASCL1 as a key regulator of the spatiotemporal order of glial lineage diversity in cortical GM and callosal WM and implicate ASCL1 dysregulation as an underlying mechanism in the pathogenesis of gliomas.

## INTRODUCTION

Neurogenesis in the neocortex follows an “inside-out” pattern in which neurons in lower cortical layers are generated before those in more superficial layers (Molyneaux et al., 2007). Consequently, neuronal laminar identity and diversity are determined in part by the timing of progenitor cell division. Whether a comparable spatiotemporal program governs gliogenesis, the generation of morphologically and functionally diverse astrocyte and oligodendrocyte lineages, in the cortex and corpus callosum remains poorly understood.

During CNS development, multiple signaling pathways such as fibroblast growth factor (FGF) signaling, bone morphogenetic protein (BMP) signaling, along with NFIA and SOX9 transcription factors contribute to the transition from neurogenesis to gliogenesis (Deneen et al., 2006; Bond et al., 2012; Kang et al., 2012; Dinh Duong et al., 2019). Within the developing forebrain, oligodendrocyte precursor cells (OPCs) are generated in three spatially and temporally distinct waves. The first two waves arise ventrally from NKX2.1-expressing and GSX2-expressing glial progenitors (GPs) in the preoptic area and ganglionic eminences between embryonic day (E)12.5 and E15.5, followed by a third wave from EMX1-expressing GPs in the dorsal pallium beginning around E15.5 and extending into the perinatal period (Kessaris et al., 2006; Yue et al., 2006). Sonic Hedgehog (SHH), secreted from the ventral forebrain and later from migrating GABAergic interneurons, is essential for inducing OLIG2 expression, a basic helix-loop-helix (bHLH) transcription factor indispensable for OPC fate specification (Sussel et al., 1999; Lu et al., 2002; Winkler et al., 2018). In parallel, the proneural factor ASCL1 also contributes to OPC specification and proliferation, including in the postnatal CNS (Parras et al., 2007; Kelenis et al., 2018). By contrast, astrocyte precursor cells (APCs) emerge from regionally patterned radial glia, reflecting spatially restricted programs of astrocyte production and diversification (Tsai et al., 2012). NOTCH signaling, acting through RBPJ, downstream HES transcriptional repressors, and NFIA, is critical for specifying APC over OPC fate while maintaining the progenitor pools (Ge et al., 2002; Namihira et al., 2009; Imayoshi et al., 2010; Kang et al., 2012).

Despite these advances, the transcriptional programs implicated in the specification and development of astrocytes or oligodendrocytes are broadly expressed in GPs and not restricted to just a single glial lineage. For instance, ASCL1 and OLIG2 are expressed in the majority of ventral GPs in the ganglionic eminences and, to a lesser extent, in GPs of the dorsal pallium (Parras et al., 2002; Miyoshi et al., 2007). Lineage-tracing and functional studies further demonstrate that ASCL1 and OLIG2 are important for both oligodendrocyte and astrocyte development (Marshall et al., 2005; Masahira et al., 2006; Yue et al., 2006; Cai et al., 2007; Vue et al., 2014). Similarly, NOTCH signaling is essential for the specification and differentiation of both APCs and OPCs (Ge et al., 2002; Genoud et al., 2002; Park and Appel, 2003; Zhang et al., 2009). Mechanistically, prominent NOTCH pathway genes (*Notch1, Dll1/3, Hes1*/*5*), along with *Nfia* and *Sox9*, are direct targets of ASCL1 and OLIG2, while HES proteins reciprocally repress the expression of these proneural factors (Kageyama et al., 2005; Castro et al., 2011; Yu et al., 2013; Myers et al., 2024). Thus, within this shared and highly interconnected transcriptional network, precisely how GPs are specified to adopt distinct APC versus OPC fates in gray matter (GM) and white matter (WM) remains unresolved.

To address this question, we characterized the temporal and spatial dynamics of ASCL1 and OLIG2 expression and tested their functional roles during gliogenesis in the developing dorsal forebrain. We show that ASCL1 and OLIG2 co-expression dynamics drive the spatiotemporal order of gliogenesis, which proceeds from the upper cortex inward toward the corpus callosum. Moreover, ASCL1 plays a primary role in promoting the migration and proliferation of GPs out of the ventricular zone (VZ) to the overlying nascent corpus callosum and cortical plate, whereas the dosage of OLIG2, induced by ASCL1, determines APC versus OPC fate decisions. Finally, we demonstrate that sustained ASCL1 expression is critical for maintaining OPCs into the postnatal brain, and ASCL1 needs to be downregulated for OPCs to differentiate into mature oligodendrocytes. Together, our results reveal that the expression dynamics of ASCL1, acting in concert with OLIG2, serve as a central spatiotemporal framework that links glial cell fate specification and differentiation with GM and WM regional identity in the dorsal forebrain.

## MATERIALS and METHODS

### Mouse Strains

The following transgenic mouse strains were used and have been previously described: *Ascl1^CreER^*knock-in [Ascl1^tm1.1(Cre/ERT2)Jejo^/J] (Kim et al., 2011); *R26R-tdTom* [Rosa26^LSL-tdTomato^] (Madisen et al., 2010); *Ascl1-floxed* [Ascl1^flox/flox^] (Pacary et al., 2011); *Olig2-floxed* [Olig2^flox/flox^] (Yue et al., 2006); *R26R-tTA* [Rosa26^LSL-tTA^] (Wang et al., 2008); *TetO-AIG* [TetO^Ascl1-ires-GFP^] (Ueki et al., 2015). Genotypes for all strains were validated using TransnetYX commercial genotyping service with either custom-designed or publicly available primers. All animal procedures were performed in accordance with NIH guidelines and were approved by the Institutional Animal Care and Use Committee (IACUC) at the University of New Mexico Health Science Center.

### Tamoxifen Administration

The presence of a vaginal plug was considered E0.5, and the day of birth was designated at as postnatal day 0 (P0). Tamoxifen (T5648, Sigma-Aldrich Co., St. Louis, MO 63103, United States) was dissolved in a 10% ethanol/90% sunflower oil mix. A single dose of tamoxifen (2.5 mg per 40 g body weight) was administered via intraperitoneal injection into pregnant females at E17.5 or subcutaneously to the back of neonatal pups at P0 or P2. Because tamoxifen administration at late gestation increased risk of birth complications, pregnant dams injected at E17.5 underwent cesarean section at E19, and pups were carefully introduced to a foster dam and raised until the designated collection time point.

### Plasmid Injection and Electroporation of Neonatal Mouse Pups

Neonatal electroporation was performed as previously described (Myers et al., 2024). Briefly, approximately 1uL of *FUGW-Cre* plasmids [1ug/uL] were injected into the lateral ventricle of P0 pups using a Picospritzer III. Plasmids were electroporated (5 pulses of 100 V, 50 ms duration, 950 ms interval) into radial glia lining the right lateral ventricle using a NEPA21 Super Electroporator and CUY650-P5 electrokinetic forceps. Each pup underwent two rounds of electroporation separated by approximately 5 minutes to enhance plasmid uptake and recombination efficiency, thereby increasing labeling of radial glia and their progeny with tdTomato or GFP.

### Mouse Perfusion, Tissue Preparation, and Immunofluorescence

Mice of both sexes were used for all analyses. Embryonic and neonatal mice were anesthetized by brief hypothermia on ice, whereas adult mice were anesthetized with Avertin (2.5 g 2,2,2-tribromoethanol dissolved in 5 mL 2-methyl-2-butanol and diluted in 200 mL distilled water).

Cardiac perfusion was performed using an Ismatec peristaltic pump with phosphate-buffered saline (PBS) for 5 minutes, followed by 4% paraformaldehyde (PFA) for 7 minutes. Brains were dissected and post-fixed overnight in 4% PFA at 4°C, washed twice in PBS (4 hours per wash), and cryoprotected in 30% sucrose. Tissue was embedded in O.C.T. compound (Fisher Healthcare) and sectioned coronally at 40 µm using a cryostat. Mouse spinal cords were sectioned at 10 µm directly onto Superfrost Plus™ slides.

For immunofluorescence, free-floating brain sections or spinal cord sections on slides were washed twice in PBS and blocked for 1 hour at room temperature in PBS containing 2% donkey serum and 0.3% Triton X-100. Sections were incubated overnight at 4°C with primary antibodies diluted in blocking solution. The following day, sections were washed three times in PBS (10 minutes each) and incubated with species-appropriate Alexa Fluor-conjugated secondary antibodies (488, 568, and/or 647; Thermo Fisher Scientific). Sections were counterstained with DAPI (10 µg/mL in PBS) for 30 minutes at room temperature, mounted onto Superfrost Plus™ slides, and coverslipped with Fluoromount-G mounting medium (SouthernBiotech). The following antibodies were used in this study:

| **Antibodies** | **Source** | **IHC Concentration** |
| --- | --- | --- |
| Chicken anti-GFP | Aveslabs / GFP-1010 | 1:1000 |
| Goat anti-OLIG2 | R&D Systems / AF2418 | 1:1000 |
| Goat anti-SOX10 | R&D Systems / AF2864 | 1:100 |
| Mouse anti-APC, clone CC-1 | Millipore Sigma / MABC200 | 1:100 |
| Rabbit anti-ASCL1 | Abcam / AB211327 | 1:1000 |
| Rabbit anti-GFAP | Millipore Sigma / AB5804 | 1:500 |
| Rabbit anti-OLIG2 | Millipore Sigma / AB9610 | 1:1000 |
| Rat anti-PDGFRα | BD Pharmingen / 558774 | 1:100 |
| Rat anti-SOX2 | ThermoFisher / Btjce | 1:1000 |

### Image Acquisition and Co-localization Quantification Analyses of Cells

Whole-brain images were acquired using a ZEISS Axioscan slide scanner (20x air objective) at the UNM Preclinical Core. High-magnification immunofluorescent images for co-localization analyses were acquired using a Leica TCS SP8 laser scanning confocal microscope (Leica Microsystems) equipped with 405, 488, 568, and 647 nm solid-state laser lines and high-sensitivity hybrid photodetectors (HyD) in UNM Comprehensive Cancer Center Microscopy Share Resources facility.

To ensure the validity of quantitative co-localization comparisons, all brain sections for a given experiment were processed for immunofluorescence imaging using identical conditions and parameters. For cell counting and co-localization analyses of various markers in the dorsal forebrain, automated tile scans were acquired using a 20x HC PL APO 0.75NA CS2 oil-immersion objective. The Leica Application Suite X (LAS X) software was used to define the scan area and automatically stitch the tiles, with a 10% overlap between adjacent tiles to ensure seamless merging. Images were captured at a resolution of 1024×1024 pixels with a scan speed of 400 Hz and a line averaging of 4 to improve the signal-to-noise ratio. The confocal pinhole was set to 1 Airy Unit (AU). For each brain, a minimum of three non-overlapping fields of view were imaged from 3-5 coronal sections for each region of interest (e.g., dorsal cortex, corpus callosum) at level of the primary motor and somatosensory areas. Raw Leica Image Files (.lif) were then imported into Imaris software (v. 9.8, Oxford Instruments) for 3D visualization, processing, and quantitative analysis.

#### Image Pre-processing

To correct for minor variations in background fluorescence and uneven illumination, a baseline subtraction was first applied to each channel using a Gaussian filter with a filter width of 10 µm, a value larger than the objects of interest (cell nuclei).

#### Automated Cell Detection and Counting

Individual fluorescently labeled cells were identified and counted using the ‘Spots’ detection module in Imaris. This algorithm models objects as spherical spots based on their fluorescence intensity profile. The estimated XY diameter for spot detection was set to 6 µm, based on empirical measurements of DAPI-stained nuclei in the tissue of neonatal mice. The ‘Background Subtraction’ option within the Spots creation wizard was enabled to enhance the signal-to-noise ratio before detection. A quality-based threshold was then applied to the resulting spots to filter out low-intensity noise. This threshold was determined empirically for each antibody channel on representative control images and was then held constant for the analysis of all images in that experiment. The total number of spots (cells) was automatically calculated by the software.

#### Definition of Regions of Interest (ROIs) and Co-localization Analysis

Anatomical regions of interest (e.g., ventricular zone, subventricular zone, intermediate zone, cortical plate, corpus callosum, upper cortical layers 1-4, lower cortical layers 5/6) were manually delineated on each image based on cytoarchitecture revealed by the DAPI counterstaining. The number of spots within each defined ROI was exported for density calculations. To determine the proportion of cells expressing multiple markers (e.g., tdTOM+/ASCL1+), the ‘Colocalize Spots’ function in Imaris was utilized. A maximum distance threshold of 5 µm between the centers of spots from different channels was set as the criterion for colocalization. The software then generated a new population of spots representing only the colocalized cells, and the counts were exported for analysis. Cell densities were calculated by dividing the total cell count within an ROI by the area of that ROI (in mm²), which was measured using Imaris.

#### Quantification of Labelled Cells in the Spinal Cord

Quantification of the number and distribution of tdTOM+ and GFP+ cells in P14 spinal cord was performed as previously reported (Vue et al., 2014).

#### Quantification of Immunofluorescence Intensity

Cellular immunofluorescence intensity for ASCL1, OLIG2, and SOX2 was measured from high-magnification images (TIFF) of triple immunostained brain or spinal cord sections as previously described (Kelenis et al., 2018). Briefly, ImageJ was used to draw outline of cells expressing markers of interest, then the area, integrated density, and mean fluorescence intensity were measured. Multiple mean background fluorescence around measured cells were also calculated. The total corrected cellular fluorescence (TCCF) was calculated using the formula: TCCF = integrated density − (cell area × mean background fluorescence) and expressed in arbitrary units to determine the cellular levels of ASCL1, OLIG2, and SOX2. Comparison of TCCF of ASCL1 in *Ascl1*-CE GFP⁺ cells with TCCF of endogenous ASCL1 and OLIG2 at E16.5 and P14 were obtained from 10 μm spinal cord sections that were processed simultaneously using identical antibody and imaging parameters.

### Statistical Analyses

Quantitative data were graphed and analyzed using GraphPad Prism software (v. 9.0, GraphPad Software, San Diego, CA). Data are presented as the mean ± standard error of the mean (SEM). For analyses involving cell counts, densities, or percentage distribution from tissue sections, N=3-4 hemi brain sections/stage, which included both males and females across at least two litters, were used for control and experimental genotypes. Quantifications were obtained for cortex, corpus callosum from 3-5 coronal sections at the level of the primary motor and somatosensory areas per brain, which were then averaged to obtain a single value for each biological replicate. Unpaired, two-tailed Student’s t-test was used to determine statistical significance between two experimental groups or conditions. A p-value of less than 0.05 was considered statistically significant.

## RESULTS

### Glial progenitors co-express ASCL1 and OLIG2 in the SVZ and IZ in the developing dorsal forebrain

To gain insights into gliogenesis in the developing dorsal forebrain, we first characterize the spatiotemporal expression dynamics of key transcription factors, ASCL1, OLIG2, and SOX2, from the ventricular zone (VZ) to the cortical plate at E17.5, P0, and P2. At these stages, neurogenesis in the cortex is mostly complete, and cells in the VZ are GPs that predominantly give rise to astrocyte and oligodendrocyte lineages (Levers et al., 2001).

Triple immunofluorescence staining for ASCL1, OLIG2, and SOX2, along with DAPI, on coronal brain sections showed that SOX2 is a global marker of gliogenesis in the dorsal pallium. Specifically, SOX2 is highly expressed in all GPs in the VZ and subventricular zone (SVZ) at all stages examined. SOX2 is also detected in GPs in the intermediate zone (IZ), which is the nascent corpus callosum, and in glial cells that have reached the cortical plate (Fig. 1Dii-Fii), but at much lower levels compared to in the VZ/SVZ (Fig. 1Gi-Ii, J-L).

**Figure 1.**
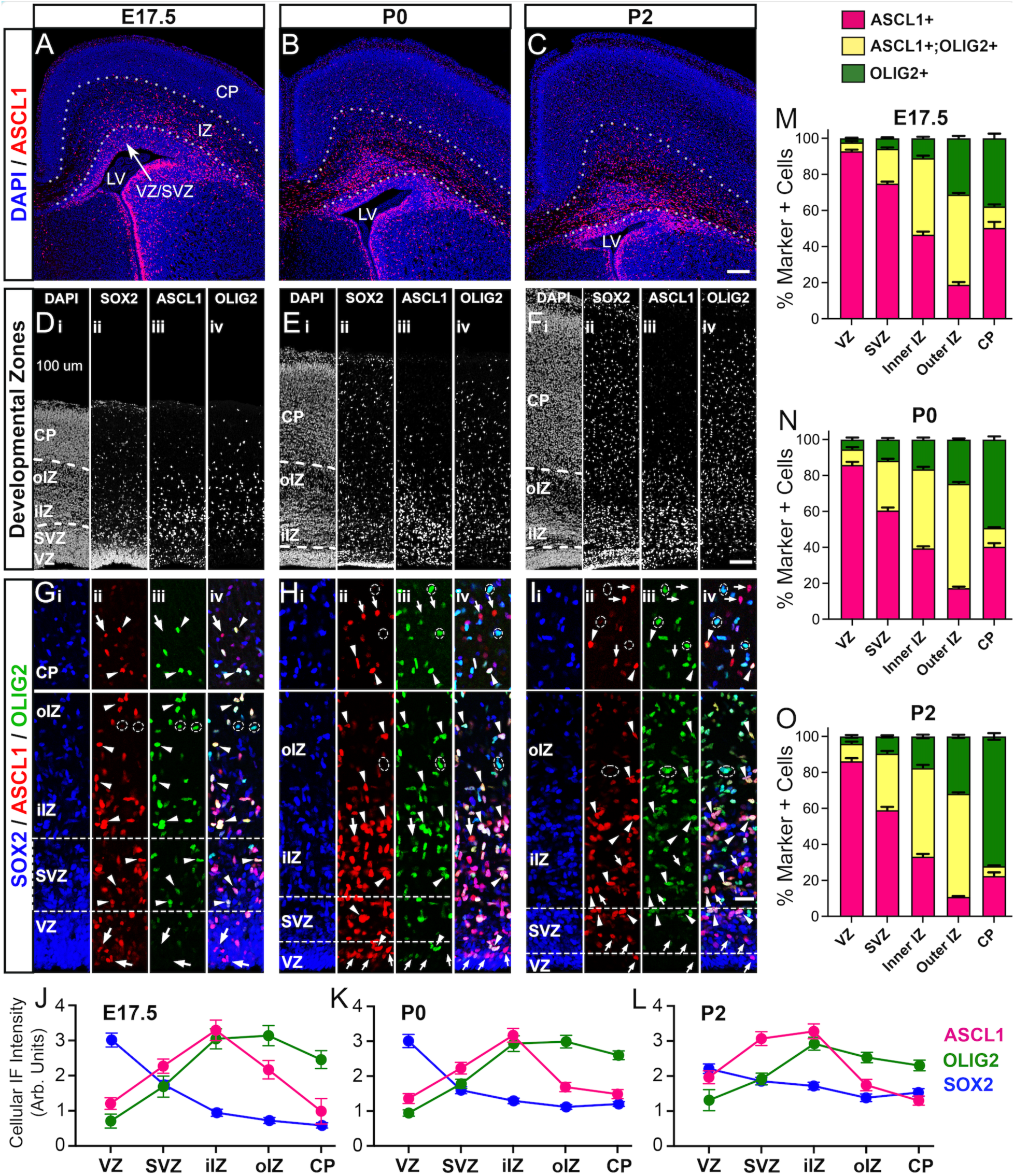
ASCL1 and OLIG2 co-expression dynamics are increased as glial progenitors transition from the ventricular zone into the intermediate zone. **(A-C)** ASCL1 immunofluorescence with DAPI on E17.5, P0, and P2 coronal brain sections showing VZ/SVZ, intermediate zone (IZ), and cortical plate (CP). **(D-F)** SOX2, ASCL1, OLIG2, and DAPI staining highlighting the cortical developmental zones. Highly dense SOX2 delineates VZ from SVZ, both of which decreases in thickness from E17.5 to P2 (Dii, Eii, Fii). ASCL1 is sparsely expressed in VZ, increased in SVZ and IZ, and downregulated in CP (Diii, Eiii, Fiii). OLIG2 starts in SVZ then maintained through IZ to CP (Div, Eiv, Fiv). **(G-I)** High magnification images showing ASCL1 expression precedes OLIG2 in the VZ (arrows), both transcription factors highly co-expressed in the SVZ and IZ and to some extent in CP (arrowheads). Dashed ovals highlight OLIG2-only cells in the outer IZ and CP. Arrows in CP highlight ASCL1-only cells. **(J-L)** Quantification of cellular levels of ASCL1, OLIG2, and SOX2 in GPs in the cortical developmental zones at E17.5 (J), P0 (K), and P2 (L). Data was obtained from N=2 brains/stage, ∼200 cells/brain at levels shown in D-F. **(M-O)** Proportion of ASCL1+ only, ASCL1+;OLIG2+ double-positive, and OLIG2+ only cells in the developmental zones at E17.5, P0, and P2. Data was obtained from N=3 brains/stage across two litters, with 3-5 sections/brain. Scale bar = 150 μm for A-C; 100 μm for D-F; 25 μm for G-I.

Unlike SOX2, ASCL1 is only expressed in a subset of GPs in the VZ, then prominently increased in the majority of SOX2+ GPs in the SVZ and IZ (Fig. 1A-C, Gii-Iii). ASCL1 expression then diminished from the outer IZ to the cortical plate and was restricted to only a subset of the SOX2+ cells (Fig. 1Diii-Fiii). Overall, ASCL1 expression follows a transient pattern that is initiated in the VZ, peaked in the inner IZ, and starts to be downregulated as glial cells exit the outer IZ and enter the cortical plate (Fig. 1J-L).

Lastly, OLIG2 expression is sparsely detected in a few GPs starting in the SVZ then increases in the IZ and continues to be maintained at high levels as glial cells entered the entire cortical plate, including those in the outermost layers at P0 and P2 (Fig. 1Div-Fiv, Giii-Iiii). Markedly, the levels of OLIG2 from the VZ to the cortical plate are inversely correlated with SOX2 (Fig. 1J-L). Consistent with previous findings that *Olig2* is a transcriptional target of ASCL1 (Vue et al., 2020), OLIG2 is highly co-expressed with ASCL1 in GPs in the SVZ and IZ at all stages examined (arrowheads, Fig. 1Giii-Iiii). However, in the cortical plate many OLIG2+ cells expressed very little or no ASCL1, especially at P0 and P2 (dotted circles, Fig. 1Giii-Iiii). Similarly, the few ASCL1+ cells in the cortical plate also do not co-express OLIG2 (arrows, top panels, Fig. 1G-I).

By quantifying the proportions of cells that were only ASCL1+, both ASCL1+ and OLIG2+, or only OLIG2+ in the various developmental zones, we found that that these cell populations vary within each developmental zone but were quite similar across the three developmental stages (Fig. 1M-O). Notably, from E17.5 to P2, the majority of GPs in the VZ (80-90%) and SVZ (60-70%) were only ASCL1+ cells (pink bar), likely because they have yet to express OLIG2. The proportion of ASCL1+ only cells then decreased in the inner and outer IZ, then increased again in the cortical plate. In contrast, the percentage of ASCL1+;OLIG2+ cells (yellow bar) dramatically increased from 5-10% in the VZ to 50-60% in the outer IZ, then decreased to 5-10% in the cortical plate.

The percentage of OLIG2+ only cells (green bar), however, increased from 5-10% in the VZ/SVZ to 40-50% in the IZ and as much as 75% in the cortical plate across the three developmental stages. Interestingly, within the cortical plate, glial cells seemed to be either ASCL1+ only or OLIG2+ only cells, but not both (Fig. 1M-O), highlighting a divergence in glial lineages.

Collectively, these findings illustrate that gliogenesis in the dorsal pallium is reliably marked by SOX2 from the VZ to the cortical plate. Furthermore, GPs first express ASCL1 in the VZ, followed by co-expression OLIG2 as the levels of ASCL1 increases and they migrate through the IZ to eventually become astrocyte or oligodendrocyte lineages that would later occupy the cortex and corpus callosum.

### ASCL1 expression is rapidly downregulated in glial progenitors in the VZ/SVZ but is sustained in the IZ

To better characterize the temporal dynamics of ASCL1 expression from the VZ to the CP, we next perform short-term lineage tracing of ASCL1-expressing GPs by crossing *Ascl1^CreRT2/+^* knock-in heterozygous mice with Cre-dependent tdTomato reporter (*R26R-tdTOM*) mice. Because *CreER* expression is under the control of the endogenous *Ascl1* promoter, tdTOM expression following tamoxifen administration faithfully labels ASCL1-lineage cells in the CNS (Kim et al., 2011). Tamoxifen was administered to *Ascl1^CreRT2/+^;R26R-tdTOM* mice at E17.5, P0, or P2 for analysis 2 days post-administration (at P0, P2, and P4, respectively) to characterize the dynamics of ASCL1 and OLIG2 expression in tdTOM-labeled cells.

In all tamoxifen stages, analysis of coronal brain sections showed that tdTOM+ cells were primarily found concentrated in the SVZ and IZ (Fig. 2A-C, D-F), in similar pattern to ASCL1 from 2 days earlier (Fig. 1A-C). Interestingly, the majority of tdTOM+ cells in the SVZ (∼70%) and inner IZ (∼50%) were negative for ASCL1 and also did not express OLIG2 (Fig. 2D-F; arrows Fig. 2J-L; red bar graph, Fig. 2M-O). This suggests that ASCL1 expression from the VZ to the inner IZ is transitory and must have been downregulated in most of the tdTOM+ cells labeled in these regions. In contrast, about 70-80% of tdTOM+ cells in the outer IZ not only continued to express ASCL1 but also OLIG2 (arrowheads, Fig. 2G-I; blue bar graph, Fig. 2M-O), highlighting that the high ASCL1 expression in the inner IZ (Fig. 1J-L) is likely maintained from 2 days earlier, which resulted in the induction of OLIG2 in tdTOM+ cells. In support of this interpretation, we found that the gradual increase in the proportion of tdTOM+ cells that were ASCL1+ from the SVZ to the inner and outer IZ closely mirrored the proportion that were OLIG2+ across all stages examined (Fig. 2P,Q). Intriguingly, for the few tdTOM+ cells that have reached the cortical plate, 30-50% were OLIG2+ only (yellow bar), followed by some that were either positive (blue bar, 20-30%) or negative (red bar, 20-30%) for both ASCL1 and OLIG2, while there were hardly any tdTOM+ cells that were ASCL1+ only (pink bar, 0-10%) (Fig. 2M-O). Indeed, across all tamoxifen stages examined, approximately 70% of tdTOM+ cells in the cortical plate were OLIG2+ while about 20-40% were ASCL1+, which further demonstrates that ASCL1 is rapidly downregulated as glial cells enter the cortical plate (Fig. 2M-O).

**Figure 2.**
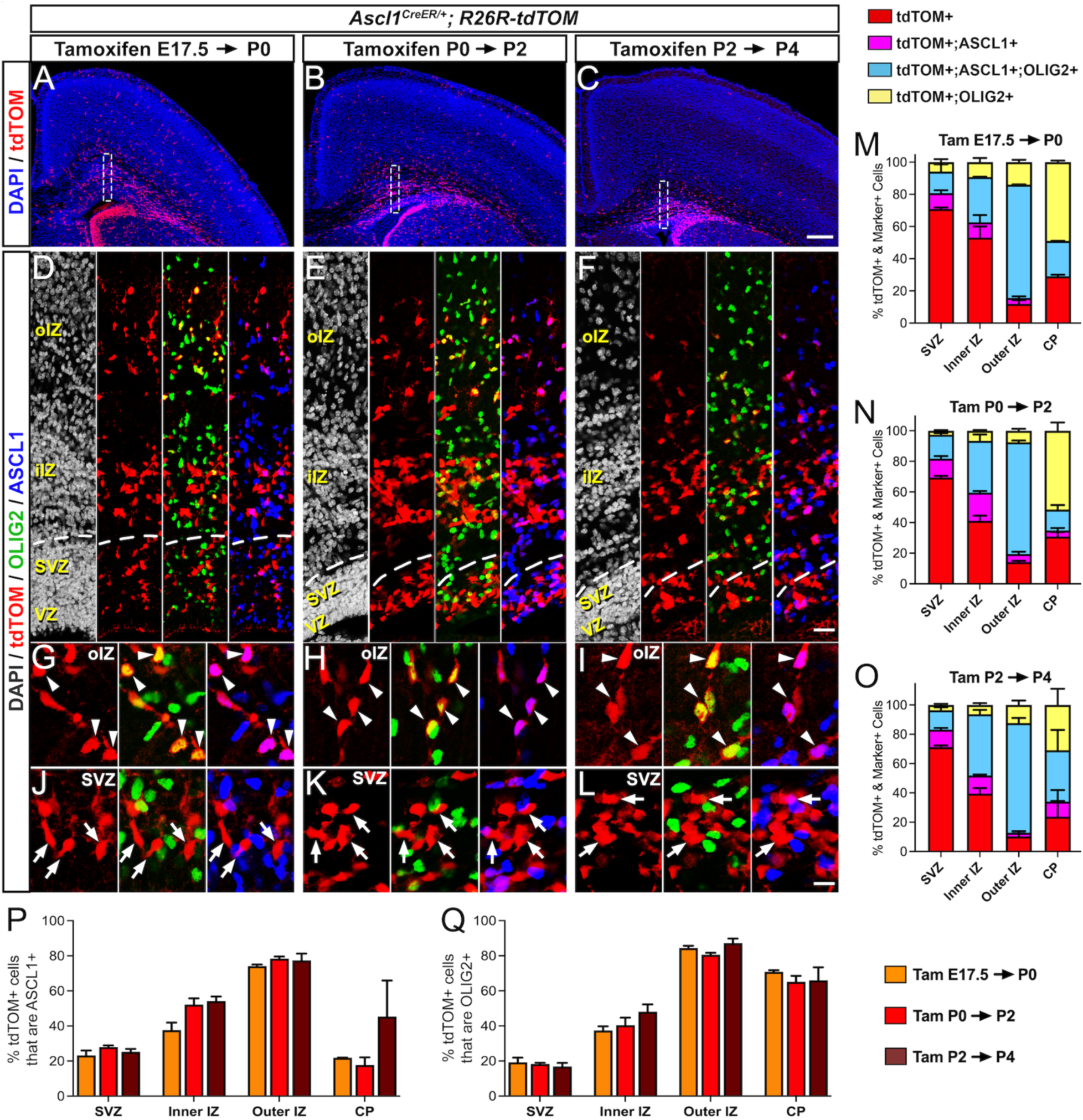
ASCL1 expression is transitory in the ventricular zone, sustained in the intermediate zone, and downregulated in the cortical plate. **(A-C**) tdTOM and DAPI staining on coronal sections of *Ascl1^CreER/+^;R26R-tdTOM* mice injected with tamoxifen at E17.5 (A), P0 (B), or P2 (C) for short-term lineage of ASCL1+ GPs. (**D-L**) Higher magnification of insets in A-C showing tdTOM, ASCL1, and OLIG2 expression in VZ, SVZ, and inner and outer IZ. tdTOM+ cells do not express ASCL1 or OLIG2 in the SVZ (arrows, J-L) but they do in the outer IZ (arrowheads, G-I). (**M-O**) Proportion of tdTOM+ cells that do or do not co-express ASCL1 and/or OLIG2 from SVZ to cortical plate. (**P,Q**) Proportion of tdTOM+ cells that are ASCL1+ (P) or OLIG2+ (Q) across the developmental zones in the three stages of tamoxifen injection. Quantification was obtained from N=3 brains/stage from two litters, with 3-5 sections/brain. Scale bar = 250 μm for A-C; 25 μm for D-F; 12.5 μm for G-L.

Taken together, because ASCL1 expression precedes OLIG2 and GPs showed a gradual increase in the co-expression of ASCL1 and OLIG2 as they migrate from the VZ/SVZ to the outer IZ, these findings indicate that the co-expression of these two bHLH transcription factors are likely responsible for driving gliogenesis in the dorsal forebrain.

### Gliogenesis follows an “outside-in” spatiotemporal pattern in the dorsal cortex and corpus callosum

Classic birth dating studies using tritiated thymidine or BrdU show that neurogenesis in the cortex follows an “inside-out” pattern, with neurons in deeper cortical layers being generated before those in more superficial layers (Angevine and Sidman, 1961; Levers et al., 2001). Unfortunately, because GPs are first specified into highly proliferative precursor cells (APCs and OPCs) before becoming postmitotic astrocytes and oligodendrocytes, such labeling method is not reliable to associate the “birthdates” of GPs with the spatial distribution of their astrocyte or oligodendrocyte progenies in the cortex and corpus callosum (Levers et al., 2001). To overcome this caveat, we took advantage of the transient nature of ASCL1 expression, which increase from VZ/SVZ to the inner IZ but is downregulated once the cells entered the cortical plate (Figs. 1&2). By administering tamoxifen at E17.5, P0, or P2 to *Ascl1^CreRT2/+^;R26R-tdTOM* mice, we predict that we should be able to, in principle, label or “birthdate” the lineages of temporally distinct GP populations with tdTOM.

As predicted, analysis of tdTOM+ cells on coronal sections of P30 brains revealed a striking “outside-in” pattern of gliogenesis (Fig. 3A-C), a stark contrast to the “inside-out” pattern of neurogenesis. Specifically, labeling of ASCL1+ GPs with tamoxifen administration at E17.5 showed that tdTOM+ progenies were largely confined to upper cortical layers 1-4, while very few were found in lower layers 5/6, and almost no cells at all were seen in the corpus callosum (Fig. 3D). On the other hand, tdTOM+ cells labeled at P0 were found across all cortical layers and showed an increased in abundance in the corpus callosum (Fig. 3E). Lastly, tdTOM+ cells labeled at P2 were mostly relegated to lower layers 5/6 and especially the corpus callosum (Fig. 3F).

**Figure 3.**
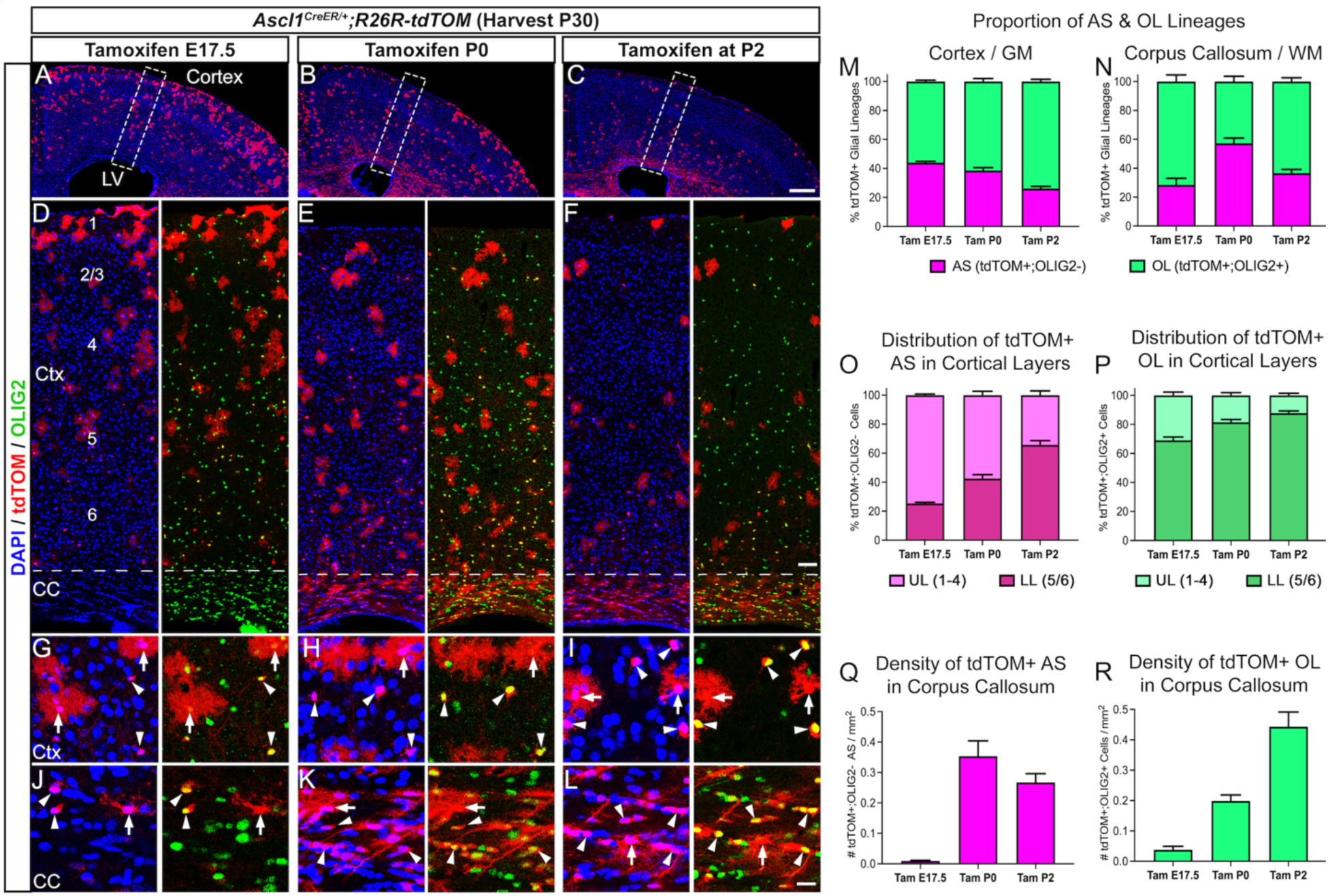
ASCL1-expressing glial progenitors give rise to both astrocyte and oligodendrocyte lineages in an “outside-in” pattern in the neocortex and corpus callosum. (**A-C**) tdTOM and DAPI staining of coronal brain sections from P30 *Ascl1^CreER/+^;R26R-tdTOM* mice injected with tamoxifen at E17.5 (A), P0 (B), or P2 (C) showing distribution of ASCL1+ GP progenies in the dorsal forebrain. (**D-F**) High magnification of insets in A-C showing tdTOM, OLIG2, and DAPI staining of cortex and corpus callosum (CC). tdTOM+ cells labeled at E17.5 occuppied predominantly upper cortical layers 1-4 while those labeled at P0 and P2 occupied lower cortical layers 5/6 and CC. (**G-L**) High magnification images showing tdTOM+ cells in cortex and CC are comprised of AS (arrows) and OL (arrowheads) based on morphology and co-localization with OLIG2. (**M,N**) Proportion of tdTOM-labeled cells that are of AS or OL lineages in Ctx (M) and CC (N). (**O,P**) Proportion of tdTOM+ AS and OL lineages in the upper and lower cortical layers. (**Q,R**) Density of tdTOM+ AS and OL lineages in CC. Quantification was obtained from N=3 brains/stage, with 3-5 sections/brain at the level of the motor and somatosensory cortex. Scale bar = 300 μm for A-C; 50 μm for D-F; 12.5 μm for G-L.

Morphological analysis combined with lineage marker immunostaining revealed that tdTOM+ cells from all three tamoxifen stages comprised of both astrocyte (arrows, OLIG2-) and oligodendrocyte (arrowheads, OLIG2+) lineages whether in the cortex or corpus callosum at P30 (Fig. 3G-L), which was similar to that observed in the spinal cord (Vue et al., 2014). However, the relative proportions of tdTOM+ astrocytes and oligodendrocytes varied depending on the stage labeled. For example, in the cortex, tamoxifen at E17.5 and P0 resulted in similar proportions of tdTOM+ astrocytes (∼40%) and oligodendrocytes (∼60%), but these proportions were shifted to ∼25% astrocytes and ∼75% oligodendrocytes with labeling at P2 (Fig. 3M). Within the corpus callosum, tamoxifen at E17.5 labeled very few tdTOM+ cells, with ∼30% being astrocytes and ∼70% being oligodendrocytes. However, this was reversed with labeling at P0, where ∼60% were astrocytes and ∼40% were oligodendrocytes. The proportions of astrocytes and oligodendrocytes then reversed again with tamoxifen at P2, in which ∼35% of tdTOM+ cells were astrocytes and ∼65% were oligodendrocytes (Fig. 3N).

We next examined the laminar distribution and density of tdTOM+ astrocyte or oligodendrocyte lineages in the cortex and corpus callosum, respectively. tdTOM+ astrocytes labeled at E17.5 were found mostly in the upper cortical layers 1-4 (∼80%). However, this pattern shifted inward and was reversed with labeling at P2, where the majority of tdTOM+ astrocytes now residing in the lower cortical layers 5/6 (∼70%) (Fig. 3O). Unlike astrocytes, tdTOM+ oligodendrocytes were consistently more concentrated in the lower cortical layers 5/6 (∼70–90%) across all labeling timepoints (Fig. 3P). This suggests that while astrocytes in all cortical layers are likely generated from E17.5 to P2, oligodendrocytes in the upper cortical layers may be generated from dorsal GPs prior to E17.5 or potentially from the earlier ventral waves of OPCs that were not labeled in this study.

In the corpus callosum, the density of tdTOM+ cells deriving from ASCL1+ GPs at E17.5 was very low, resulting in only a few oligodendrocytes and virtually no astrocytes labeled. However, the density of tdTOM+ cells labeled at P0 and P2 increased markedly for both lineages (Fig. 3Q). Notably, the density of tdTOM+ oligodendrocytes showed a 2-fold increased between each successive labeling timepoint (Fig. 3R), highlighting the robust expansion of dorsally-derived OPCs within the first postnatal week.

Taken together, these results reveal that gliogenesis in the dorsal pallium follows an “outside-in” spatiotemporal pattern where cortical GM astrocytes and oligodendrocytes are likely generated embryonically until birth, while WM astrocytes and oligodendrocytes in the corpus callosum are primarily generated around and after birth. Thus, astrocyte and oligodendrocyte lineages in the cortex and corpus callosum may be derived from temporally and/or spatially distinct GP populations.

### Constitutive expression of *Ascl1* specifies and maintains OPCs in the postnatal brain

Lineage tracing consistently showed that ASCL1+ GPs highly co-express OLIG2 and give rise to both astrocyte and oligodendrocyte lineages in the brain and spinal cord (Fig. 3) (Vue et al., 2014). This then begs the questions, how are GPs specified to adopt astrocyte versus oligodendrocyte fates, and what roles do ASCL1 and OLIG2 play in this process?

Since ASCL1 expression precedes OLIG2 in GPs in the VZ, but OLIG2 is essential for OPC fate specification (Lu et al., 2002), we hypothesized that sustained ASCL1 levels are required to induce sufficient OLIG2 levels to specify GPs as OPCs rather than APCs. To test this, we crossed *Ascl1^CreRT2/+^* knock-in allele into mice carrying dual alleles of a Cre-dependent tetracycline transactivator (*R26R-tTA*) and a TetO-promoter driving expression of *Ascl1-ires-GFP* bicistronic transcript. The combination of all three alleles allows for induction of constitutive expression of *Ascl1* (*Ascl1*-CE) and *GFP* in ASCL1+ GPs following tamoxifen administration (Fig. 4A). We first administered tamoxifen at E14.5 to *Ascl1^CreRT2/+^;R26R-Ascl1*-CE embryos to measure the immunofluorescent levels of ASCL1 in GFP+ cells at P14, given its continuous expression once induced. We found that ASCL1 levels in GFP+ cells were similar to endogenous ASCL1 levels at E16.5, which were significantly higher than the endogenous ASCL1 levels at P14 (Fig. 4B). This indicates that ASCL1 levels driven by *Ascl1*-CE are within biologically relevant levels appropriate to address its role in glial cell fate specification.

**Figure 4.**
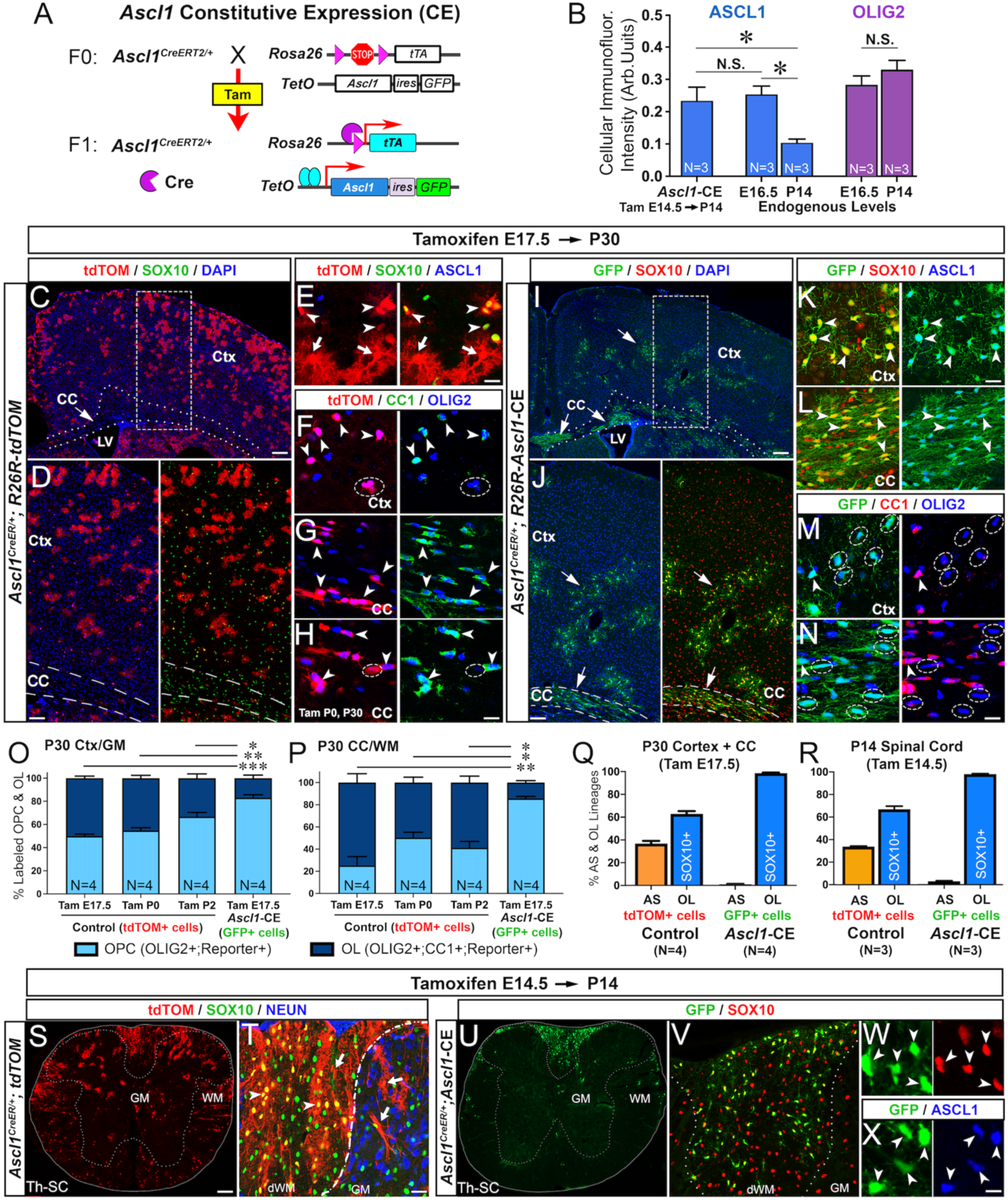
Constitutive ASCL1 expression specifies glial progenitors exclusively as OPCs in the brain and spinal cord. **(A)** Schematic of transgenic alleles and mouse breeding used to induce *Ascl1* constitutive expression (CE), along with *GFP*, in GPs following tamoxifen injection. **(B)** Quantification of the cellular levels of ASCL1 based on immunofluorescent intensity in *Ascl1-*CE GFP+ cells compared to normal cells in E16.5 and P14 spinal cord sections. Note endogenous ASCL1 levels significantly decreased from E16.5 to P14 but OLIG2 levels do not, which serve as internal control. (**C-H**) Coronal brain sections from P30 *Ascl1^CreER/+^;R26R-tdTOM* control mice tamoxifen at E17.5 or P0. tdTOM+ cells are mostly in upper Ctx (C,D), no longer express ASCL1 (E), and comprised of both AS (arrows) and OL (arrowheads, SOX10+, OLIG2+) lineages, many of which have differentiated into CC1+ OLs (E-H). (**I-N**) Coronal brain sections from P30 *Ascl1^CreER/+^;R26R-Ascl1-*CE mice tamoxifen at E17.5. GFP+ cells are found as clusters mainly in lower Ctx and CC, comprised exclusively of OL (arrowheads, SOX10+, OLIG2+) lineage that are negative for CC1 (dotted ovals, M,N). (**O,P**) Proportion of OLIG2+;tdTOM+ or OLIG2+;GFP+ cells derived from the various stages of tamoxifen (E17.5 to P2) that are CC1+ (OLs) or CC1-(OPCs) in Ctx/GM or CC/WM at P30. (**Q,R**) Proportion of tdTOM+ or GFP+ cells that are AS (SOX10-, orange) or OL (SOX10+, blue) lineages in brain (Q) or spinal cord (R). (**S-X**) Transverse thoracic spinal cord sections of P14 *Ascl1^CreER/+^;R26R-tdTOM* control (O,P) and *Ascl1^CreER/+^;R26R-Ascl1-CE* (Q-S) mice (tamoxifen at E14.5) showing tdTOM+ and GFP+ cells in the gray matter (GM) and white matter (WM). tdTOM+ cells are comprised of both AS (arrows, SOX10-) and OL (arrowheads, SOX10+) lineages (P,S), but GFP+ cells are only OL lineage (arrowheads, SOX10+). Scale bar = 200 μm for C, I; 100 μm for D, J, S, U; 25 μm for T, V; 12.5 μm for E-H, K-N, W, X.

We then administered tamoxifen at E17.5 to *Ascl1^CreRT2/+^;R26R-Ascl1*-CE mice for direct comparison with *Ascl1^CreRT2/+^;R26-tdTOM* control mice at P30 in the brain. Unlike tdTOM+ control cells, which were found distributed mostly in the upper cortex and comprised of ∼40% astrocyte (arrows, SOX10-) and ∼60% oligodendrocyte (arrowheads, SOX10+) lineage cells (Fig. 4C-E,Q), GFP+ cells were found as clusters mainly in the lower cortex as well as in the corpus callosum (arrows, Fig. 4I,J) and were comprised of 100% SOX10+ oligodendrocyte lineages (arrowheads, Fig. 4K-N,Q). Interestingly, the morphology and clustering of the GFP+ cells were very different from SOX10+;tdTOM+ cells, indicating that they could be proliferating OPCs rather than oligodendrocytes. To confirm their identity, immunostaining revealed that most of the OLIG2+;GFP+ cells in the cortex and corpus callosum did not express the oligodendrocyte marker, CC1 (dotted circles, Fig. 4M,N). This was unlike the OLIG2+; tdTOM+ control cells, which were mostly CC1+ oligodendrocytes (arrowheads, Fig. 4F-H). Specifically, ∼50% of OLIG2+;tdTOM+ cells in the cortex and ∼75% in the corpus callosum were CC1+, while less than 20% of OLIG2+;GFP+ cells in both regions were CC1+ (first & last bar graphs, Fig. 4O,P). This decrease in the rate of differentiation of OLIG2+;GFP+ cells into CC1+ oligodendrocytes was also significant when compared to OLIG2+;tdTOM+ cells that were labeled at P0 or P2 (middle bar graphs, Fig. 4O,P).

To determine if the sustained ASCL1 levels also specify OPC fate in other regions of the CNS, we administered tamoxifen at E14.5 to label ASCL1+ GPs during gliogenesis in the spinal cord of *Ascl1^CreRT2/+^;R26R-Ascl1*-CE mice for direct comparison with *Ascl1^CreRT2/+^;R26R-tdTOM* controls. As seen in the brain, we found that at P14, while tdTOM+ cells in control spinal cords were consisted of ∼35% astrocyte (arrows) and ∼65% oligodendrocytes (arrowheads, SOX10+) lineage cells (Fig. 4R-T), GFP+ cells were exclusively SOX10+ oligodendrocyte lineage cells (Fig. 4R,U-X).

Taken together, our results revealed that sustained developmental levels of ASCL1 are sufficient to specify GPs into oligodendroglial lineage while also maintains them as OPCs into the postnatal CNS, likely by suppressing their differentiation into postmitotic CC1+ oligodendrocytes.

### ASCL1 promotes glial progenitor proliferation and migration but requires OLIG2 for specification and survival of OPCs

Next, we aimed to determine whether induction of *Ascl1*-CE was sufficient to specify OPC fate from radial glia, which are quiescent neural stem cells lining the ventricle that normally express very low or no ASCL1. To investigate this, we injected then electroporated a Cre-expressing plasmid (*FUGW-Cre*) into the right SVZ of *R26R-Ascl1*-CE mice at birth for both short-term (P6) and long-term (P30) lineage tracing analyses in comparison to *R26R-tdTOM* control mice (Fig. 5A-C).

**Figure 5.**
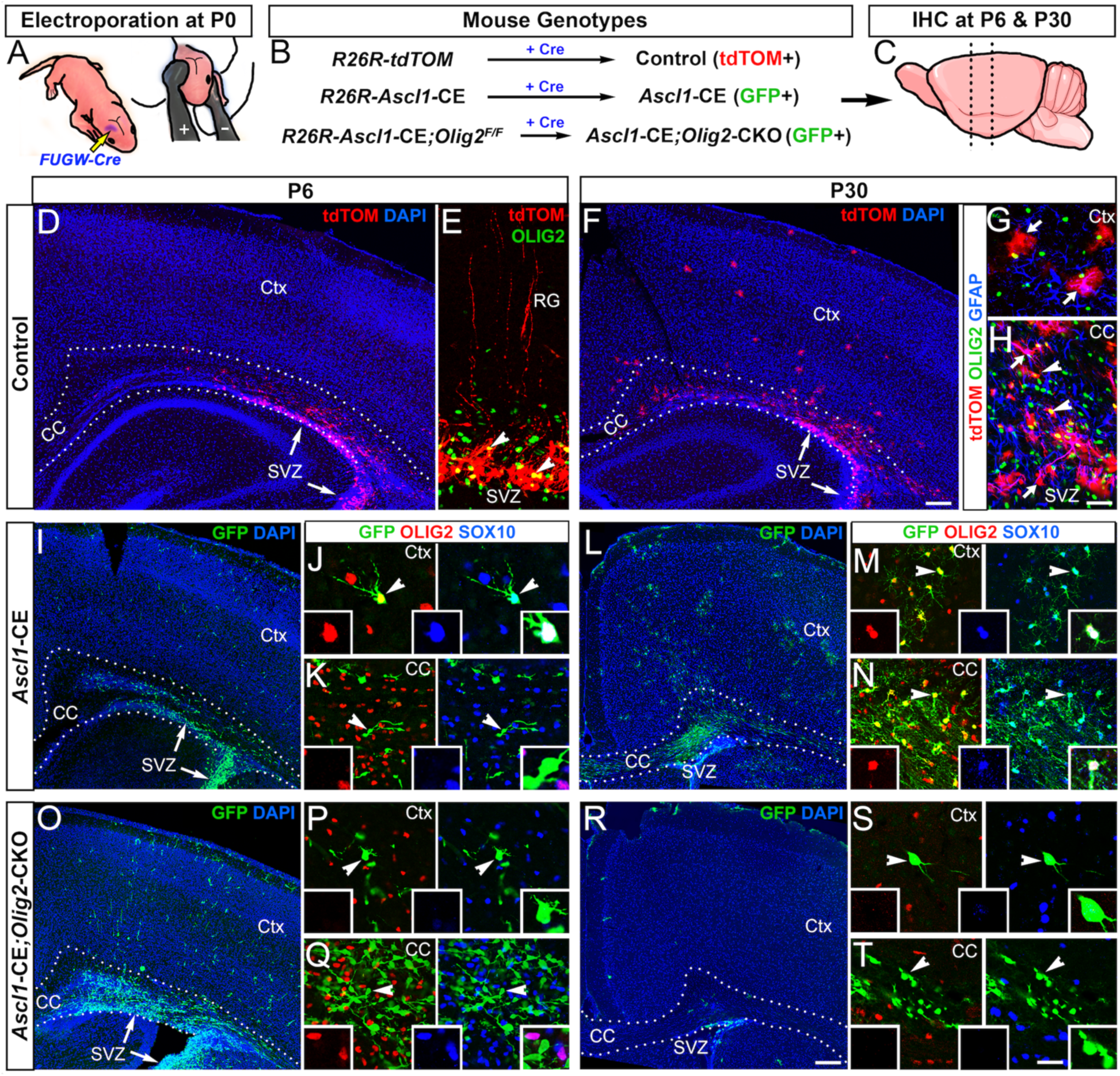
ASCL1 independently promotes proliferation and migration of glial progenitors but requires OLIG2 to specify and maintain the survival of OPCs. **(A-C)** Schematic of injection and electroporation of *FUGW-Cre* plasmids into neural progenitors in the SVZ of P0 mouse pups with the genotypes indicated for analysis at P6 and P30. (**D-H**) Staining of coronal brain sections of control mice showing electroporated tdTOM+ cells at P6 and P30. At P6, tdTOM+ cells are radial glia (RG) in SVZ with a few expressing OLIG2 (arrowheads, D,E). At P30, tdTOM+ cells are found in SVZ, corpus callosum, and cortex, and are comprised of both AS (arrows) and OL (arrowheads, OLIG2+) lineage cells. (**I-N**) Staining of coronal brain sections of *Ascl1-*CE mice showing GFP+ cells at P6 and P30. Number of GFP+ cells is increased in the SVZ and CC as well as in Ctx at P6. OLIG2 and SOX10 are not expressed in GFP+ cells in SVZ and CC but are expressed in some GFP+ cells in Ctx (arrowheads and insets, I,J). At P30, GFP+ cells formed clusters in both Ctx and CC (K) and are exclusively OL (OLIG2+,SOX10+) lineages (arrowheads, insets, L,M). (**O-T**) Staining of coronal brain sections of *Ascl1-*CE*;Olig2-*CKO mice showing GFP+ cells at P6 and P30. Number of GFP+ cells is dramatically increased in SVZ, CC, and Ctx at P6 (even compared to *Ascl1-CE*). Note that SOX10 is not expressed in GFP+ cells in Ctx (arrowheads and insets, O,P). At P30, GFP+ cells do not express OLIG2 or SOX10 and are reduced numbers in Ctx and CC. Phenotype was observed in 4-6 brains/genotype. Scale bar = 200 μm for D, F, I, L, O, R; 25 μm for the rest and 12.5 μm for insets.

Analysis of coronal brain sections at P6 showed that electroporated tdTOM+ cells were found at SVZ surface, many of which possess long radial processes extending through the cortex (Fig. 5D,E). Immunostaining revealed that only a few of these tdTOM+ cells were OLIG2+ (arrowheads, Fig. 5E). By P30, some tdTOM+ cells have migrated to the cortex and gave rise to predominantly protoplasmic astrocytes (arrows), based on morphology and co-expression with GFAP but not with OLIG2 (Fig. 5F,G). The majority of tdTOM+ cells, however, were found still in the SVZ and corpus callosum, and while some were OLIG2+ (arrowheads) or GFAP+ (arrows), most were negative for these markers (Fig. 5F,H).

In contrast, induction of *Ascl1*-CE showed that there was a dramatic increase in the number of GFP+ cells in the SVZ, corpus callosum, and the cortex at P6, including in the superficial marginal zone, highlighting an increase in both cell proliferation and migration (Fig. 5I). Unlike the tdTOM+ control cells, some of the GFP+ cells in the cortex already co-expressed both OLIG2 and SOX10, indicating that they were oligodendrocyte lineage cells (Fig. 5J). Interestingly, the GFP+ cells in the SVZ and corpus callosum have yet to express OLIG2 or SOX10 (Fig. 5K). By P30, the GFP+ cells in both the cortex and corpus callosum were all OLIG2+ and SOX10+ (Fig. 5L-N). The GFP+ cells at this stage also formed large clusters, especially in the corpus callosum, indicating that they are proliferating OPCs that failed to differentiate to become postmitotic CC1+ oligodendrocytes.

Given that OLIG2 is a direct target of ASCL1 and these two bHLH transcription factors can physically interact to bind to DNA (Myers et al., 2024), we next asked which aspect of ASCL1’s ability to promote GP proliferation, migration, and OPC fate specification is dependent on OLIG2. To accomplish this, we electroporated *FUGW-Cre* plasmids into the right SVZ to simultaneously induce *Ascl1*-CE while conditionally knocking out *Olig2* (*Ascl1*-CE;*Olig2*-CKO) in radial glia (Fig. 5B). Surprisingly, we observed an even greater number of GFP+ cells in the SVZ, corpus callosum, and cortex (Fig. 5O) when compared to *Ascl1*-CE alone at P6 (Fig. 5I). This suggest that ASCL1’s impact on cell proliferation and migration were enhanced in the absence of OLIG2. Without OLIG2, SOX10 was not expressed in any of the GFP+ cells that had migrated into the cortex at P6 (Fig. 5P,O). By P30, unexpectedly the *Ascl1*-CE;*Olig2*-CKO GFP+ cells had mostly disappeared from the cortex and corpus callosum, except for a few cells that remained at SVZ surface, corpus callosum, and in the marginal zone of the cortex (Fig. 5R). These remaining GFP+ cells were still negative for SOX10 and exhibited aberrant morphologies that seemed to be neither astrocyte or oligodendrocyte in identity (arrowheads, Fig. 5S,T).

Collectively, these findings demonstrate that ASCL1 induces GP proliferation and migration from the VZ to the cortex but acts through and requires OLIG2 to specify and preserve the survival of OPCs (SOX10+) into the postnatal CNS. It is possible that ASCL1 expression also needs to be downregulated to properly specified APCs and maintained the survival of astrocyte lineage cells since *Ascl1*-CE;*Olig2*-CKO GFP+ cells vanished by P30.

### Chronic ASCL1 expression maintains OPCs in an undifferentiated glioma-like state

ASCL1’s ability to promote cell proliferation and migration while suppressing differentiation of OPCs under constitutive context suggests that it may possess oncogenic potential. To assess this, *Ascl1*-CE mice (N=5) electroporated with *FUGW-Cre* plasmids at P0 was maintained until 7 months of age. Although none of the mice exhibited noticeable neurological symptoms, analysis of both sagittal and coronal brain sections showed large clusters of GFP+ cells that had significantly expanded in size and were widely distributed along both the rostral-caudal and medial-lateral aspects of the electroporated hemisphere, including to the olfactory bulb and thalamus (Fig. 6A,B). The density and morphology of the GFP+ cells highly expressed OPC markers OLIG2 and PDGFRA but not the astrocyte marker GFAP and resembled early-stage gliomas (Fig. 6C-J). This finding demonstrates that a sustained or constitutive expression of ASCL1 is sufficient to imbue glial progenitor or precursor cells with aberrant self-renewal capacity that could easily be co-opted to initiate gliomagenesis in the brain.

**Figure 6.**
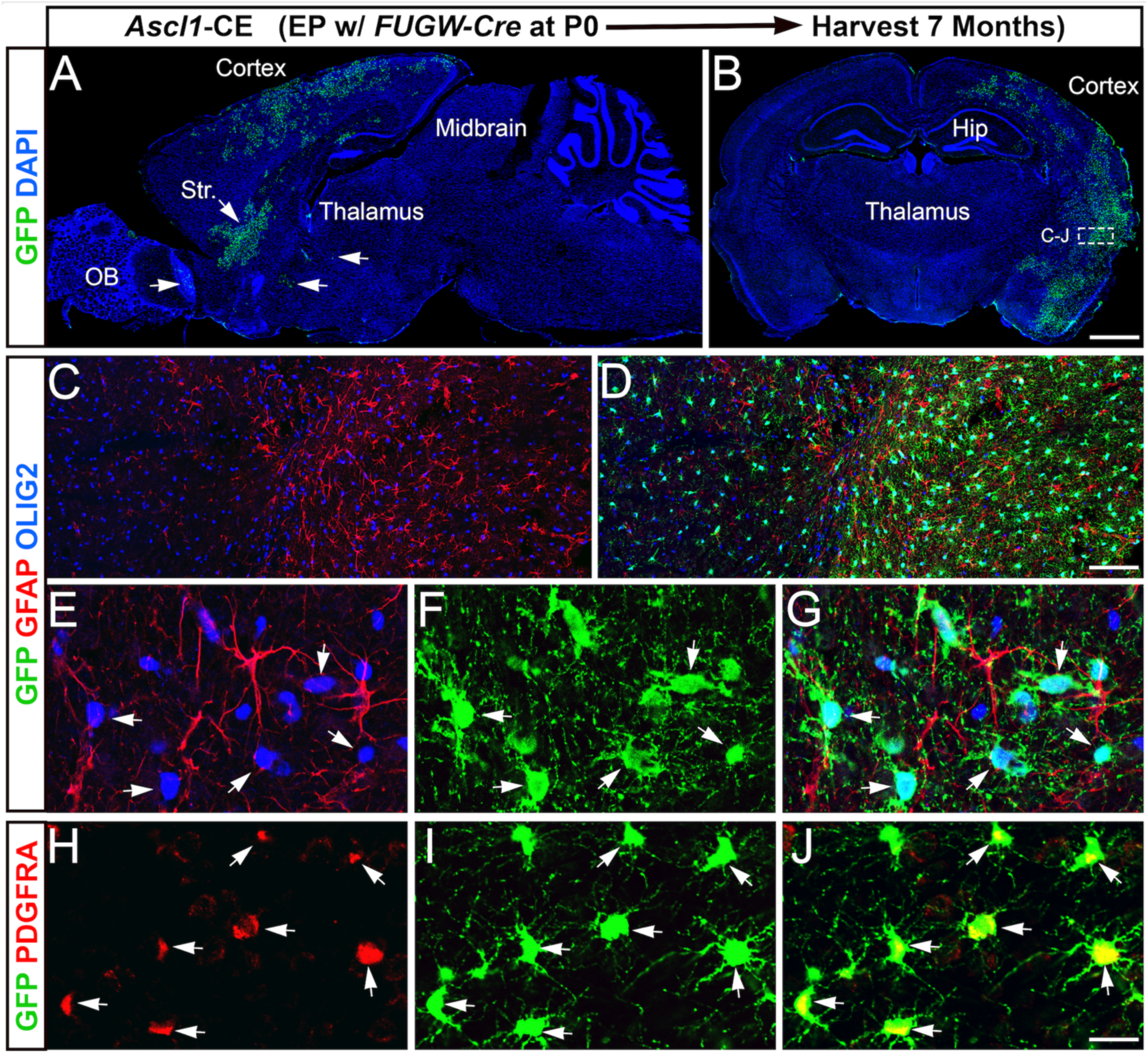
Chronic expression of ASCL1 maintains OPCs in an undifferentiated glioma-like state. **(A,B)** GFP and DAPI staining on sagittal and coronal brain sections of 7-month-old *Ascl1-*CE mice electroporated at P0. Note that GFP+ cells formed large glioma-like clusters, which are distributed throughout the electroporated hemisphere and distant brain regions (arrows, A). (**C-J**) High magnification images of GFP+ cell clusters stained for GFAP and OLIG2 (C-G) or PDGFRA (H-J). All GFP+ cells co-localize with OLIG2 and OPC marker PDGFRA (arrows, E-J) but not with astrocyte GFAP. This phenotype was observed in N=5 brains. Scale bar = 1 mm for A, B; 100 μm C, D; 20 μm for E-J.

## DISCUSSION

In this study, we show that SOX2 reliably marks glial lineages from the VZ to the cortical plate in the developing dorsal forebrain. In contrast, ASCL1 follows a dynamic trajectory, initiating sparsely in the VZ, peaking in the intermediate zone (IZ), and diminishing in the cortical plate. OLIG2 expression emerges slightly later than ASCL1, starting in the SVZ and expanding broadly from the IZ to the cortical plate (Fig. 1). Short-term lineage tracing revealed that ASCL1 expression is transitory in VZ/SVZ progenitors but becomes sustained as cells reach the IZ, where OLIG2 is concurrently induced (Fig. 2). Long-term lineage tracing further demonstrated that ASCL1+ progenitors, whether labeled at E17.5, P0, or P2, readily give rise to both astrocytes and oligodendrocytes in an “outside-in” spatiotemporal order, with those in upper cortical layers (1-4) generated earlier during embryonic stages, whereas those in deeper cortical layers (5/6) and the corpus callosum are generated around or after birth (Fig. 3). Finally, we show that constitutive expression of ASCL1 even in radial glia was sufficient to drive their transition into proliferating and migrating glial progenitors, followed by induction of OLIG2, which is absolutely required for OPC fate and survival (Fig. 5). Notably, sustained ASCL1 expression also maintains OPCs in a proliferative and undifferentiated state that fails to mature into CC1⁺ oligodendrocytes. (Fig. 4) Over time, these OPCs acquire glioma-like features and disseminate broadly throughout the brain (Fig. 6), highlighting the oncogenic-like potential of ASCL1 if persistently expressed under dysregulated state.

Together, these findings support a model in which the duration and amount of ASCL1 act as a temporal gate in controlling glial progenitor transition and lineage decisions in the developing forebrain (Fig. 7). For example, brief ASCL1 expression in the VZ likely drive generation of gliogenic progenitors from radial glia, whereas sustained ASCL1 expression promotes the migration and proliferation of these progenitors from the VZ into the developing corpus callosum or intermediate zone. During this transition, ASCL1 simultaneously activates *Olig2*. Once expressed, OLIG2 likely reinforces its own expression and that of *Ascl1*, as OLIG2 directly binds to cis-regulatory regions of the *Ascl1* and *Olig1/2* loci (Myers et al., 2024). This reciprocal feedforward relationship between ASCL1 and OLIG2 is consistent with their extensive co-expression in the developing spinal cord to both the ventral and dorsal forebrain (Nakatani et al., 2013; Vue et al., 2014). Eventually, glial progenitors that express sufficient levels of ASCL1 and OLIG2 are thereby programmed toward OPC fate, which in the presence of ASCL1, continue to proliferate and are maintained into postnatal stages (Figs. 4&5) (Marshall et al., 2005; Kelenis et al., 2018). In parallel, NOTCH signaling and HES-family repressors, whether activated via lateral inhibition or environmental cues, likely dampen ASCL1 and/or OLIG2 expression in neighboring glial progenitors, thereby biasing these cells toward APC fate (Namihira et al., 2009; Imayoshi et al., 2013). The spatial context of the intermediate zone with densely packed unmyelinated callosal axons may also serve as a physical checkpoint to ensure that GP fate choices are fully consolidated by regulating the levels of ASCL1 and OLIG2 via signaling pathways (Fig. 1J-L). Overall, gliogenesis is a lengthy but tightly regulated process that requires precise control of ASCL1 and OLIG2 expression dynamics to ensure proper specification and differentiation of both astrocyte and oligodendrocyte lineages.

**Figure 7.**
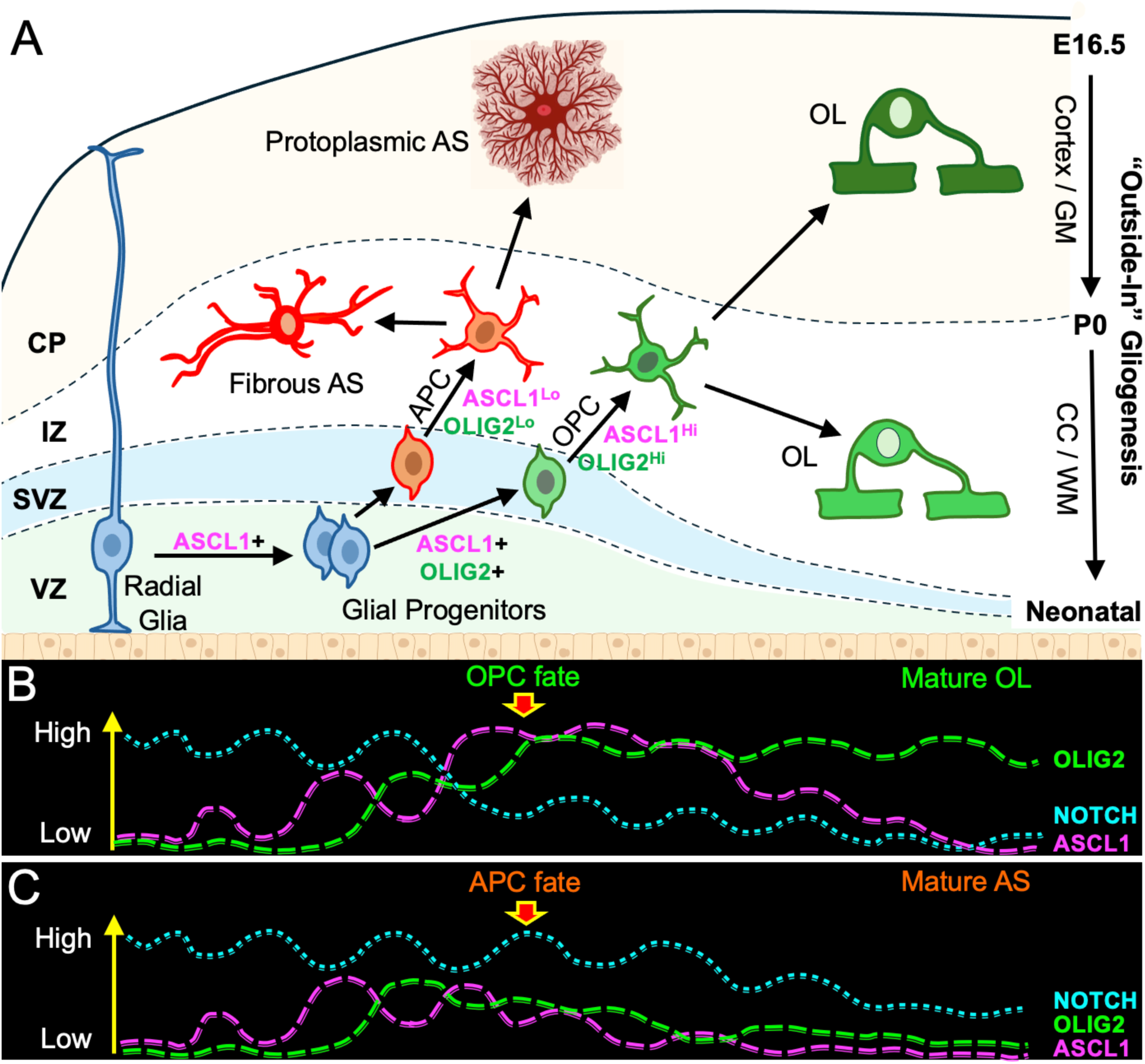
Model of spatiotemporal control of glial cell fate and regional diversity in the dorsal forebrain. **(A)** Schematic of “outside-in” pattern of gliogenesis in the cortex and corpus callosum, which starts around E16.5 to neonatal stages. ASCL1 expression promotes transition of radial glia into proliferating and migrating glial progenitors while also induces OLIG2 expression. (**B,C**) Proposed model of ASCL1 and OLIG2 co-expression dynamics and NOTCH tone in specifying glial progenitor cell fate. From SVZ to IZ, glial progenitors with high and sustained levels of ASCL1 and OLIG2 but low NOTCH tone are specified as OPCs (B), whereas glial progenitors with high NOTCH but low or transitory levels of ASCL1 and OLIG2 are specified as APCs (C). Within OPCs, downregulation of ASCL1 and NOTCH enables differentiation into oligodendrocytes, while downregulation of ASCL1 and OLIG2 in APCs leads to differentiation into astrocytes.

Unlike neurogenesis, our lineage tracing of ASCL1+ progenitors from E17.5 to P2 reveals that gliogenesis in the dorsal forebrain proceeds in an “outside-in” manner. Several intrinsic properties of glial lineages likely contribute to this pattern. Both APCs and OPCs are known to establish non-overlapping territorial domains or tiles within gray matter and white matter. For example, protoplasmic astrocytes in the cortex and hippocampus occupy discrete domains with minimal overlap, which are maintained through homotypic repulsion and contact-mediated boundary formation (Bushong et al., 2002). Similarly, OPCs are uniformly distributed in a grid-like pattern throughout the CNS and engage in mutual repulsion to prevent clustering while ensuring even coverage of both gray matter and white matter regions (Hughes et al., 2013). Because of this self-repulsive behavior, newly generated APCs and OPCs preferentially migrate into regions unoccupied by their respective lineages, resulting in a progressive inward shift from superficial cortical layers toward deeper layers and the corpus callosum, as revealed by our temporal lineage tracing.

However, it is possible that the dynamic levels of ASCL1 and OLIG2 within APCs and OPCs may contribute to the diversification of these glial lineages between gray matter and white matter. In particular, studies have shown that white matter OPCs proliferate more robustly and differentiate more readily into oligodendrocytes than their gray matter OPC counterparts in the postnatal brain and spinal cord (Psachoulia et al., 2009). This intrinsic difference may directly be linked to the higher levels of ASCL1 in white matter OPCs compared to gray matter OPCs (Kelenis et al., 2018). Consistent with this idea, ASCL1+ progenitors labeled at E17.5 with tdTOM populate the cortex extensively but largely avoid the corpus callosum, including those that are OLIG2+;tdTOM+ (Fig. 3A,D,Q,R), potentially because ASCL1 is rapidly downregulated in these early OPCs (Fig. 1J). By contrast, constitutive ASCL1 expression at E17.5 drives robust accumulation of OLIG2+;GFP⁺ OPCs in both cortex and corpus callosum, demonstrating that increasing and sustaining the levels of ASCL1 can promote white matter occupancy by OPCs at a time when they normally should not (Fig. 4I-N). A similar requirement and sufficiency for sustained ASCL1 expression in the specification of white matter OPCs was seen in the spinal cord (Fig. 4S-X) (Vue et al., 2014). Along similar line, OLIG2 may play a parallel role in astrocyte diversification into gray matter and white matter. Though not directly tested in this study, and while OLIG2 is canonically associated with OPC fate, it is transiently expressed in APCs before being downregulated in mature astrocytes, particularly in developing gray matter astrocytes (Marshall et al., 2005; Cai et al., 2007). Although the levels of OLIG2 between developing gray matter versus white matter astrocyte lineages have yet to be directly compared, *hGFAP-Cre* deletion of *Olig2* leads to a selective decrease of GFAP+ astrocytes in the corpus callosum and a concomitant increase in GFAP+ astrocytes in the cortex, a phenotype similar to *Olig1-Cre* deletion of *Rbpj*, phenotypes which are opposite to that of control wild-type brains (Cai et al., 2007; Guo et al., 2023). These studies indicate that there may be specific requirement for OLIG2 in regulating the differentiation or proper distribution of protoplasmic and fibrous astrocytes to their respective gray matter and white matter regions.

Beyond its established role in OPC fate specification, previous studies have shown that *Ascl1* is expressed at low but detectable levels in subsets of postnatal OPCs (Zhang et al., 2014; Sueda and Kageyama, 2021). However, the functional significance of ASCL1 expression in this context, particularly in OPCs in the adult brain, has remained unclear. Our study identifies ASCL1 as a key regulator of OPC self-renewal, a property that is essential for maintaining the long-lived OPC pool required to support adaptive myelination throughout life. Specifically, we demonstrate that sustained ASCL1 expression is sufficient to maintain OPCs in an undifferentiated, progenitor-like state by suppressing their differentiation into CC1⁺ oligodendrocytes. These findings support a model in which ASCL1, in concert with NOTCH signaling, acts as a transcriptional gatekeeper in balancing OPC self-renewal and differentiation in the postnatal CNS. This interpretation is consistent with genetic studies showing that OPC-specific deletion of *Ascl1* leads to decrease proliferation while deletion of *Notch1* promotes premature differentiation into MBP+ oligodendrocytes (Kelenis et al., 2018; Zhang et al., 2009). Accordingly, since *Notch1* is a direct transcriptional target of ASCL1, downregulation of ASCL1 in OPCs may be required to release the NOTCH-inhibitory tone to permit differentiation, which was not possible with *Ascl1*-CE. Neuronal activity, glutamatergic signaling, calcium-dependent pathways, and growth factor signaling represent plausible mechanisms by which ASCL1 and/or NOTCH signaling may be sustained or attenuated to favor OPC self-renewal versus differentiation. While this property positions ASCL1 as a potentially attractive therapeutic target for enhancing OPC expansion or reprogramming for remyelination purposes, our findings caution that ASCL1’s expression and function must be precisely regulated, or it may cause aberrant self-renewal that can lead to glioma initiation and progression.

Interestingly, despite the marked increased in GFP+ cells in *Ascl1*-CE;*Olig2*-CKO mice between P0 and P6, even compared to *Ascl1*-CE manipulation alone, the near complete loss of these cells at P30 suggests that sustained ASCL1 expression in the absence of OLIG2 generates an unstable cellular state. Under normal conditions, APCs transiently express some levels of ASCL1 and OLIG2 before downregulating both factors as they mature into astrocytes. Sustained ASCL1 expression combined with loss of OLIG2 may therefore promote cell proliferation and migration, as seen at P6, but ultimately trap cells in a developmental intermediate that cannot resolve into a viable or stable glial lineage. Because these cells failed to acquire a stable astrocyte or oligodendrocyte identity, they may not be able to integrate into functional neural circuits, ultimately leading to their elimination from the brain. Unfortunately, a technical limitation of our study is the inability to constitutively express OLIG2 in radial glia in the context of ASCL1 loss, which would allow for direct assessment of whether OLIG2 alone is sufficient to specify, expand, and maintain the survival of OPCs or APCs in the absence of ASCL1, and whether these precursor cells could proliferate, migrate, or differentiate under these conditions. Future studies employing this approach will be essential to further disentangle the independent versus cooperative roles of ASCL1 and OLIG2 in glial lineage specification, maturation, and long-term maintenance.

## AUTHOR CONTRIBUTIONS

L.E.P.B and T.Y.V designed research; L.E.P.B., H.C., M.L., E.V., B.L.M., J.D., A.R., and T.Y.V performed research; L.E.P.B., H.C., and T.Y.V. analyzed data; L.E.P.B. and T.Y.V. wrote the paper.

## ACKNOWLEDGEMENT

This research was supported by NINDS R01NS121660 and NIAAA P50AA022534 NMARC Pilot 6B grants (T.Y.V.) and partially by the UNM Comprehensive Cancer Center (UNMCCC) NCI P30CA118100 grant and the UNM Center for Brain Recovery and Repair (CBRR) NIGMS P20GM109089 grant. The Fluorescence Microscopy Shared Resource Facility in the UNMCCC, which receive additional support from the State of New Mexico, and the Preclinical Core in the CBRR, were essential toward the completion of this research. We thank Dr. Richard Lu for *Olig2*-floxed mice and Dr. Masato Nakafuku for *TetO-Ascl1-ires-GFP* mice.

